# Assessing Quantitative MRI Techniques using Multimodal Comparisons

**DOI:** 10.1101/2022.02.10.479780

**Authors:** Francis Carter, Alfred Anwander, Thomás Goucha, Helyne Adamson, Angela D. Friederici, Antoine Lutti, Claudine J Gauthier, Nikolaus Weiskopf, Pierre-Louis Bazin, Christopher J Steele

## Abstract

The study of brain structure and change in neuroscience is commonly conducted using macroscopic morphological measures of the brain such as regional volume or cortical thickness, providing little insight into the microstructure and physiology of the brain. In contrast, quantitative MRI allows the monitoring of microscopic brain change non-invasively in-vivo, and provides normative values for comparisons between tissues, regions, and individuals. To support the development and common use of qMRI for cognitive neuroscience, we analysed a set of qMRI metrics (R1, R2*, Magnetization Transfer saturation, Proton Density saturation, Fractional Anisotropy, Mean Diffusivity) in 101 healthy young adults. Here we provide a comprehensive descriptive analysis of these metrics and their linear relationships to each other in grey and white matter to develop a more complete understanding of the relationship to tissue microstructure. Furthermore, we provide evidence that combinations of metrics may uncover informative gradients across the brain by showing that lower variance components of PCA may be used to identify cortical gradients otherwise hidden within individual metrics. We discuss these results within the context of microstructural and physiological neuroscience research.

## Introduction

Magnetic Resonance Imaging (MRI) gives rise to a variety of image contrasts primarily driven by differences in longitudinal and transverse relaxation rates of the MRI signal between voxels. The estimates of these relaxation rates, provided by quantitative MRI (qMRI) [Guilfoyle et al., 2003; Weiskopf et al., 2021], have been shown to correlate with microstructural properties of brain tissues such as iron or myelin concentration [Fukunaga et al., 2010; Sereno et al., 2013] and show great potential for the monitoring of microscopic brain change in-vivo [Tardif et al., 2016; Weiskopf et al., 2015]. However, the specificity of these biomarkers is limited by the combined contributions of multiple histological properties to qMRI estimates [Draganski et al., 2011; Tardif et al., 2016; Tardif et al., 2017; Weiskopf et al., 2015; Weiskopf et al., 2021]. Improving their specificity requires an understanding of how individual tissue properties are reflected in qMRI estimates. Towards this goal, the current study quantifies the specificity and overlap of the information contained within a set of quantitative MRI metrics to assist in the development of “in-vivo histology”, the mapping of MRI signals to microstructural tissue properties [Weiskopf et al., 2015; Weiskopf et al., 2021].

The assessment of microscopic brain tissue properties from in-vivo qMRI data requires a detailed understanding of how these properties impact the MRI signal [Weiskopf et al., 2015; Weiskopf et al., 2021]. Recent research includes evidence that the MR relaxation rates R1 and R2* are related to myelin and iron concentrations [Kirilina et al., 2020; Möller et al., 2019; Shams et al., 2019; Stüber et al., 2014; Trampel et al., 2019; Weiskopf et al., 2015] and that magnetization transfer (MT/MTR) measurements scale with myelin’s contribution to the macromolecular content [Mangeat et al., 2015; Manning et al., 2017; van der Weijden et al., 2021]. Myelin concentration has been shown to be the dominant contributor to multiple qMRI measures [Callaghan et al., 2015a; Mancini et al., 2020], highlighting the need for complementary qMRI measures in order to achieve a complete description of brain microstructure.

One major issue is that the signal from a single MRI voxel is potentially the result of many different combinations of molecules and their concentrations (i.e., many different molecular arrangements and concentrations can give rise to the same magnetic properties) [Draganski et al., 2011; Tardif et al., 2016; Tardif et al., 2017]. Multimodal MRI – where multiple MRI contrasts are acquired in the same participant or sample – may help address this issue. Multiple qMRI measures with different relationships to tissue properties can be combined to extract latent variables that capture shared variance. Extracted latent variables can be related to microstructural features and ground-truth molecular concentrations to determine how well they map to specific tissue properties [Borsboom et al., 2003; Filo et al., 2019; Geeraert et al., 2019]. However, this requires accurate segmentation and subsequent reduction of tissue partial voluming, which has a large but often overlooked effect on the interpretation of results [Lüsebrink et al., 2013], and is often difficult to properly implement for cortical grey matter where partial voluming with dura/CSF and WM is common at standard resolutions [Dahnke et al., 2013; Han et al., 2004; Roche and Forbes, 2014]. qMRI has other advantages, including higher reproducibility and comparability between acquisitions within and between participants than conventional structural MRI commonly used for assessing macroscopic morphological change [Caan et al., 2019; Leutritz et al., 2020; Weiskopf et al., 2015]. The advent of more advanced qMRI sequences that simultaneously capture multiple quantitative metrics has led to shorter overall acquisition times [European Society of Radiology (ESR), 2015; Weiskopf et al., 2013] and removes the need for within-subject registration between metrics. Notably, this work has led to the development of Multi-Parametric Mapping, which simultaneously captures quantitative R1, R2*, MT, and PD images [MPM; Weiskopf et al., 2013] and multi-echo MP2RAGE [T1, T2*, QSM; Caan et al., 2019; Metere et al., 2017].

While useful for acquiring multiple co-registered modalities, multiparametric methods likely contain redundancies in the information provided by each metric [Callaghan et al., 2015b; Weiskopf et al., 2015; Weiskopf et al., 2021]. Redundancies have the potential to be exploited to determine which combination of metrics accounts for the most variance in the tissue of interest (and could therefore form a basis for a minimum useful multiparametric acquisition profile) and, importantly, can then be used to map putative microstructural similarities across the brain [Weiskopf et al., 2015]. Similar to the way that microstructure exhibits differences across different tissues and regions, the relationships between metrics are also expected to vary. Previous work has explored metric-metric covariance relationships with ROI-based or segmented network approaches [Seidlitz et al., 2018; Uddin et al., 2019] and a within-ROI binning approach [Filo et al., 2019], which rely on *a priori* regional delineations and averaging that may obscure subtle variability within regions. A voxel-wise approach could provide a more nuanced and comprehensive description of these relationships and help to determine if and/or how multiparametric combinations can provide additional information not found in individual metrics [Paquola et al., 2019].

The present study provides a statistical description of the linear relationships for a set of six metrics, including four qMRI metrics from the MPM [Weiskopf et al., 2013] (R1, R2*, Magnetization Transfer saturation-MT, and Proton Density saturation - PD) and two diffusion MRI metrics (dMRI: Fractional Anisotropy - FA, and Mean Diffusivity - MD). We set out to identify potential redundancies between metrics, and additionally identify the set of normative relationships between metrics across different tissue types (sub/cortical grey matter, white matter) that can be used to develop and test hypotheses in the development of in-vivo histology. We also provide evidence that multimodal imaging holds promise in describing the microstructural drivers of qMRI contrast by applying PCA to extract and map linear latent variables within the data [Geeraert et al., 2019].

## Methods

### 2.1. Participants

Our sample consisted of 101 young healthy individuals (mean age 25.7 ± 4, range 18-34 years, 25 females) who were recruited in the area of Leipzig, Germany. All participants were right-handed, had a high-school-level education, no history of neurological or psychiatric disorders, and did not use any centrally effective medication. The study was approved by the ethics committee at the medical faculty of the University of Leipzig. All participants gave written informed consent and were paid for participation in the study.

### 2.2. MRI data acquisition and quantitative metric estimation

Quantitative multi-parametric maps (MPM, 1mm iso) and high spatial and angular resolution diffusion-weighted MRI (dMRI, 1.3mm iso) were acquired for all participants on a 3-Tesla PRISMAfit MRI system (Siemens Healthineers, Erlangen, Germany) using a standard 32-channel receive head coil. The MPM protocol consisted of three multi-echo 3D FLASH (fast low angle shot) acquisitions with predominant T1-, PD-, and MT-weighting by adapting the repetition time (TR) and the flip angle (T1w: 18.7 ms/20°, PDw and MTw: 23.7 ms/6°) following the setting previously published [Weiskopf et al., 2013]. The MTw acquisition included an additional initial 4 ms long off-resonance MT saturation pulse (flip angle 220°, 2 kHz offset) [Helms et al., 2008]. We acquired six gradient echoes (equidistant echo times, TE, from 2.2 to 14.7 ms) with alternating polarity for the T1w and the MTw volumes and eight datasets for the PDw volumes (TE from 2.2 to 19.7ms). The remaining imaging parameters were: field-of-view 256 × 240 mm, 176 slices, acceleration in phase-direction using GRAPPA 2 and in partition direction using 6/8 partial Fourier, non-selective RF excitation, high readout bandwidth = 425 Hz/pixel, RF spoiling phase increment = 50°, acquisition time ∼21 min. The acquisition was preceded by additional calibration data to correct for spatial variations in the radio-frequency transmit field (B1+) as described by Lutti and colleagues [Lutti et al., 2012]. The diffusion MRI protocol used the multi-band sequence developed at CMRR (https://www.cmrr.umn.edu/multiband) with the following parameters: 1.3 mm isotropic resolution, b-value = 1000 s/mm², 60 directions and seven images without diffusion weighting (b = 0), three repetitions to improve the SNR, TE = 75 ms, TR = 6 s, GRAPPA 2, CMRR-SMS 2, two b = 0 acquisitions with opposite phase encoding directions. The acquisition time for the dMRI protocol was 23 min.

The MPMs acquisitions were processed with the freely available hMRI toolbox (http://hmri.info, Tabelow et al., 2019) to estimate parameter maps of the longitudinal relaxation rate (R1), the effective transverse relaxation rate (R2*), the magnetization transfer (MT), the proton density (PD) for each participant. The dMRI images were corrected for motion, eddy currents, and susceptibility distortions [Andersson and Sotiropoulos, 2016], and the fractional anisotropy (FA), and mean diffusivity (MD) were computed from the diffusion tensor images using the FSL tools [Jenkinson et al., 2012]. Registration from native diffusion to higher resolution MPM space was accomplished by rigidly co-registering the b0 and R1 images within each individual, and applying the transform to FA and MD maps to bring them into R1 space (trilinear interpolation).

### 2.3. Preprocessing

After data collection and quantitative metric reconstruction, the following steps were performed to segment the brains into different tissue types (cerebral cortical GM, subcortical GM, WM), and to create surface representations of the cortex. We obtained high-quality segmentations and cortical representations by employing previously established methods [Bazin et al., 2014; Huntenburg et al., 2017; Tardif et al., 2015]. Other than the SPM skull stripping and segmentations described below, all other preprocessing was performed with the open-source neuroimaging toolbox Nighres v1.2 [Huntenburg et al., 2018].

#### 2.3.1 Segmentation

Cortical, subcortical, and WM segmentations for each participant were performed with standard GM segmentation in SPM12 on the R1 image [Ashburner and Friston, 2005] and the Nighres implementation of Multiple object Geometric Deformable Model (MGDM) segmentation using FA, R1, MT, and PD [Bazin et al., 2014; Bogovic et al., 2013]. MGDM allowed for the segmentation of the cerebral cortical and subcortical GM while preserving topology, and SPM segmentations were then combined as follows to ensure final segmentations that reduced tissue partial voluming. WM masks were defined as the intersection between SPM WM probability maps thresholded at 0.5 and the MGDM cerebral WM. GM masks were defined as the intersection between the SPM GM (probability maps thresholded at 0.95) and the MGDM labels for cerebral GM. Subcortical GM masks were defined as the intersection between the MGDM subcortical labels and the SPM GM thresholded at 0.95. The resulting masks retained artifactual R2* signal (mainly in the medial temporal lobe) which we removed by excluding voxels containing R2* values above 25 s^-1^ in GM (WM was not thresholded). We also used an MD threshold of 0.001 mm^2^s^-1^ x 10^-6^ to reduce any partial voluming with CSF in the GM masks. All masking and metric thresholds were chosen to restrict our analyses as closely as possible to tissue type (GM, WM) and optimally reduce partial voluming and signal artifacts. Subsequent analyses were performed on voxel-wise values extracted from the cortical sheet, WM, and subcortical GM in both hemispheres. We also repeated these analyses without any metric thresholds and found very similar results with identical significant effects (data not shown). A group surface co-registration step for both hemispheres was additionally used to project cortical results onto a common group surface for visualization as detailed below.

#### 2.3.2 Cortex extraction, mesh generation, and surface projection

A 3D mesh representation of the cortex was used to map and visualize our results on the cortical surface. To mitigate against the effects of partial voluming on the pial and WM surfaces, we extracted a thin sheath of the middle layer of the cortex with Nighres as follows. The cortex was first extracted with CRUISE Cortex Extraction [Han et al., 2004], generating GM-WM and GM-CSF boundary levelset images (where a levelset is the distance of each voxel from each boundary; thus representing the inner and outer cortical surfaces in voxel space). These two boundaries were then combined to extract a levelset representation of the midline of the cortical sheet. The midline levelset, used later for group co-registration, was then thresholded at a distance of 0.5 mm to generate a binary mask, and converted to a 3D mesh for display. Details on segmentation and the levelset approach for the surface generation with Nighres can be found in our previous technical publications [Huntenburg et al., 2018], and all specific commands and parameters are included as code in our github repository (https://github.com/neuralabc/paper_QuantitativeMetricComparisons).

### 2.4 Group co-registration

Cortical midline levelsets were used to perform surface-based registration in voxel space with a method similar to that developed by Tardif and colleagues [Tardif et al., 2015]. The ANTs [Avants et al., 2011; Tustison et al., 2014] multivariate template construction script was used to perform nonlinear registrations and generate a group template for both hemispheres. The levelsets representing each participant’s middle cortical ribbon were thresholded at 10 mm from the cortex and used as inputs to ANTs Multivariate Template Construction (v2, 3 iterative steps, with default linear and nonlinear registration parameters, Demons similarity metric). The final template levelset was then converted to a mesh representation for display [Huntenburg et al., 2018]. To map the results of the analysis onto a mesh representation of the group template surface, participants’ data from each metric was: 1) multiplied by a binarized midline ribbon, 2) nonlinearly transformed into the study-specific group space with the computed deformation (ANTs SyN, nearest-neighbor interpolation), and 3) a 2 mm intensity propagation was then performed to project participants data onto the final coregistered surface. In all cases, the vertex-wise median values across participants were presented for display.

### 2.5. Statistical analysis

#### 2.5.1 Metric-metric density plots

Bivariate logarithmic density plots were created to describe the relationship between metrics in GM, WM, cortex, and subcortical GM. Linear regressions were used to calculate the coefficients of determination (R-squared) for each bivariate relationship. Univariate density plots (histograms) were also generated to further describe the data.

#### 2.5.2 Dimensionality reduction

We performed model-free dimensionality reduction with Principal Component Analysis (PCA) in cortical tissue to visualize latent variables within the data. All six metrics in each of these tissues from all participants were z-scored and used as input to a PCA in scikit learn (Pedregosa et al., 2011) in Python (3.7), and all six PCs were assessed. Each participant’s data was then transformed into the PCs’ subspace for visualization, and transformed into the common group space. This allowed us to compute the median across participants and visualize our results on the cortical surface template.

## 3. Results

### 3.1 Cortical segmentation

Our segmentations combined MGDM and SPM segmentations to reduce biases caused by partial voluming of GM with WM or CSF. Visual inspection showed that our results had limited partial voluming of the cortex with CSF and WM, with little to no gyral or sulcal bias (Dahnke et al., 2013; Figure 1). As an additional confirmatory step before using this segmentation in our analyses, we also checked to ensure that our cortical maps showed high correspondence with those presented in previous literature. As expected, R1 cortical maps exhibited a gradient with peaks in primary sensory and motor regions (Figure 1), a pattern that is in excellent correspondence with what has been shown in previous literature [Marques et al., 2017; Shams et al., 2019].

**Figure 1:**
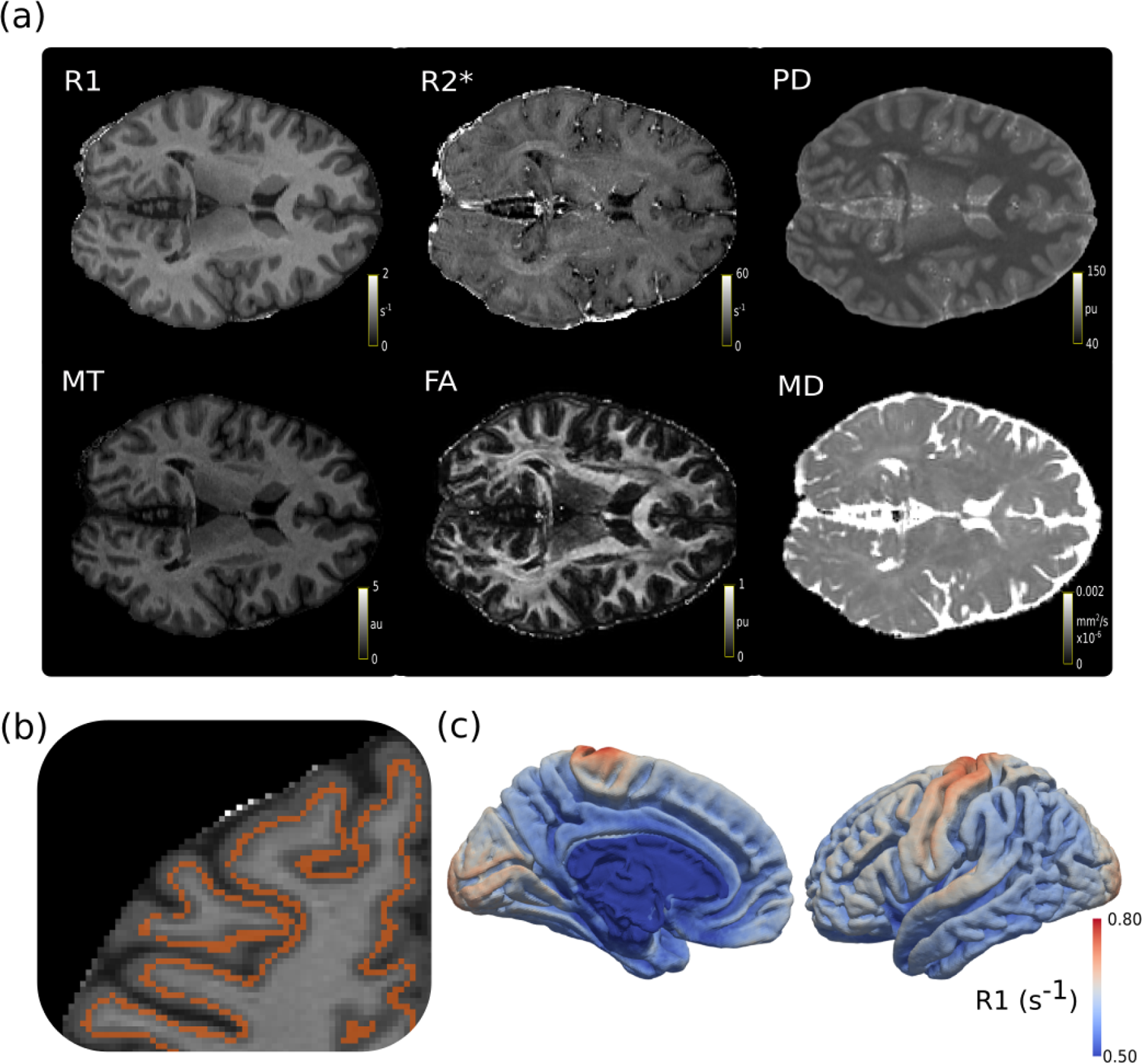
Overview of segmentation results. (a) The six metrics used were captured in 101 participants. (b) A sample brain with a red mask of the middle cortical ribbon as a semi-transparent overlay on R1, showing little to no partial voluming. (c) Using data from the cortical ribbon, we plotted the median R1 values across all participants to the group template. Other metric plots can be found in the supplemental data.

### 3.2 Descriptive statistics: GM and WM

All pairs of metrics, both in GM (combined cortical and subcortical) and WM, were found to be significantly correlated (Figure 2, all p<0.001). However, there were large fluctuations in these pairwise correlations, with R^2^ values ranging from as low as 0.0001 in the MD-R2* GM correlation, to as high as 0.482 in the R1-MT GM correlation. These differences were not equally dispersed across metrics; R1 had the highest correlations with other metrics across all structures (R^2^ average = 0.258), while MD had the weakest correlations (R^2^ average = 0.051). Furthermore, the strengths of correlations varied across tissues, with some R^2^ values differing by over 10% of explained variance. Notably, FA-MD, MD-MT, and R1-MD all had R^2^ values that were more than 10% higher in GM than in WM, and almost all pairwise correlations were greater in GM than WM. However, FA-PD and FA-R2* had R^2^ values slightly higher in WM than in GM. These relationships were reliable and consistent across participants, as shown by the low standard deviation of R^2^ across participants in Supplementary Figure 3.

**Figure 2:**
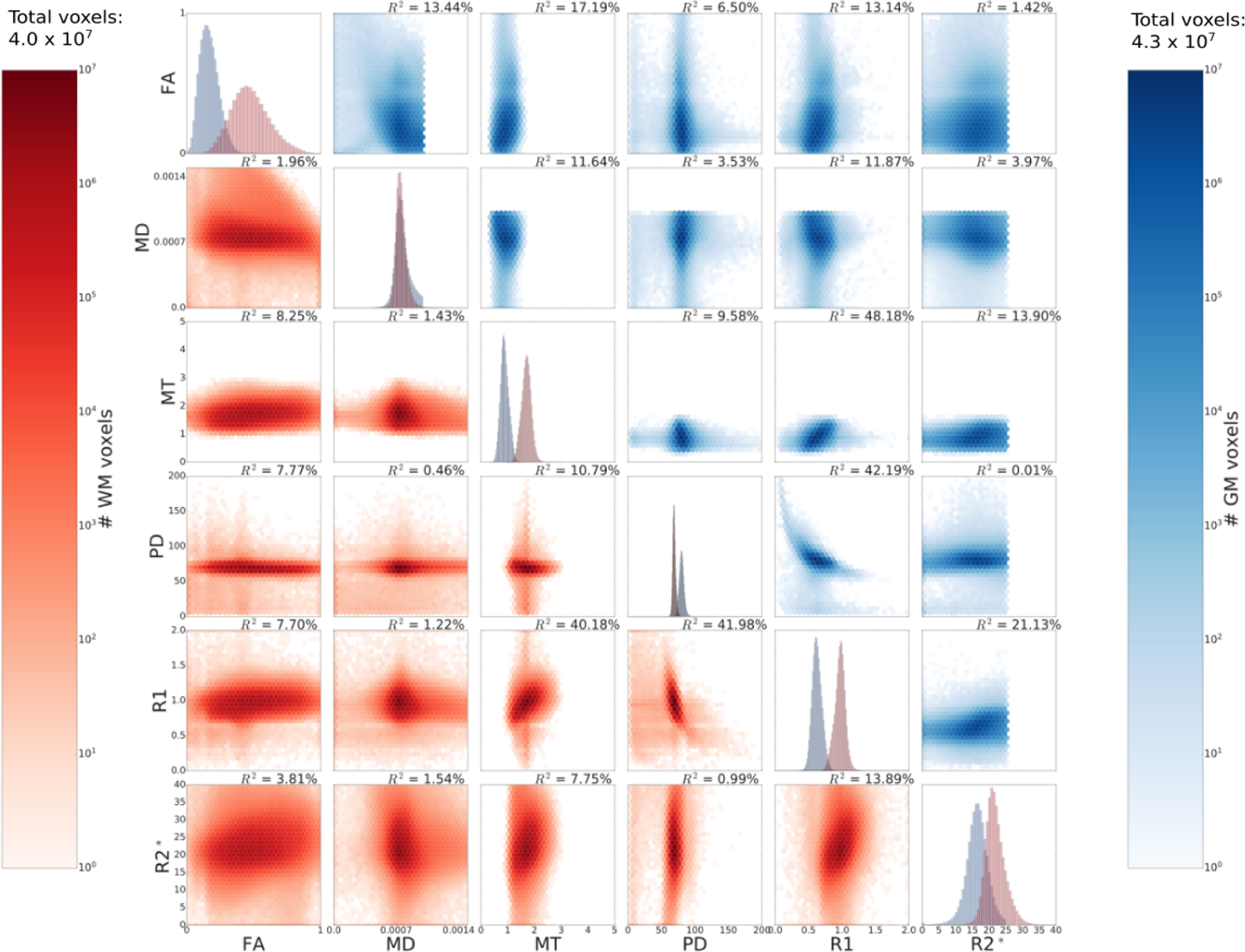
Pairwise correlations in GM (above the diagonal) and WM (below the diagonal) for all participants’ data. Data along the diagonal shows the histograms for each metric in GM (blue) and WM (red). GM includes values extracted from the midline cortical ribbon and subcortical nuclei. Metric thresholds were implemented for R2* and MD to reduce partial volume effects and artifacts, and are visible as sharp cutoffs in the figure (see Method). Pairwise density plots are depicted in log space to show the full range of data.

### 3.3 Descriptive statistics: Cortical and subcortical GM

All pairs of metrics were found to be significantly correlated in cortical and subcortical GM (Figure 3, all p<0.001). However, there were large fluctuations in these pairwise correlations, with R^2^ values ranging from as low as <0.005 in the PD-R2* cortical correlation, to as high as 0.447 in the R1-PD subcortical correlation. As with the GM and WM comparisons (Figure 2), differences were not equally dispersed across metrics; R1 had the highest correlations in all GM structures (R^2^ average = 0.230), while MD had the weakest correlations (R^2^ average = 0.071). Furthermore, the strengths of correlations varied across tissues, with some R^2^ values differing by over 10% of explained variance. Notably, FA-MD and FA-R1 correlations were 23.0 and 9.1% stronger in cortical GM than in subcortical GM, respectively. Conversely, the R1-PD and R1-R2* relationships were 11.2 and 9.1 % stronger in subcortical GM than cortical GM. These relationships were reliable and consistent across participants, as shown by the low standard deviation of R^2^ across participants in Supplementary Figure 4.

**Figure 3:**
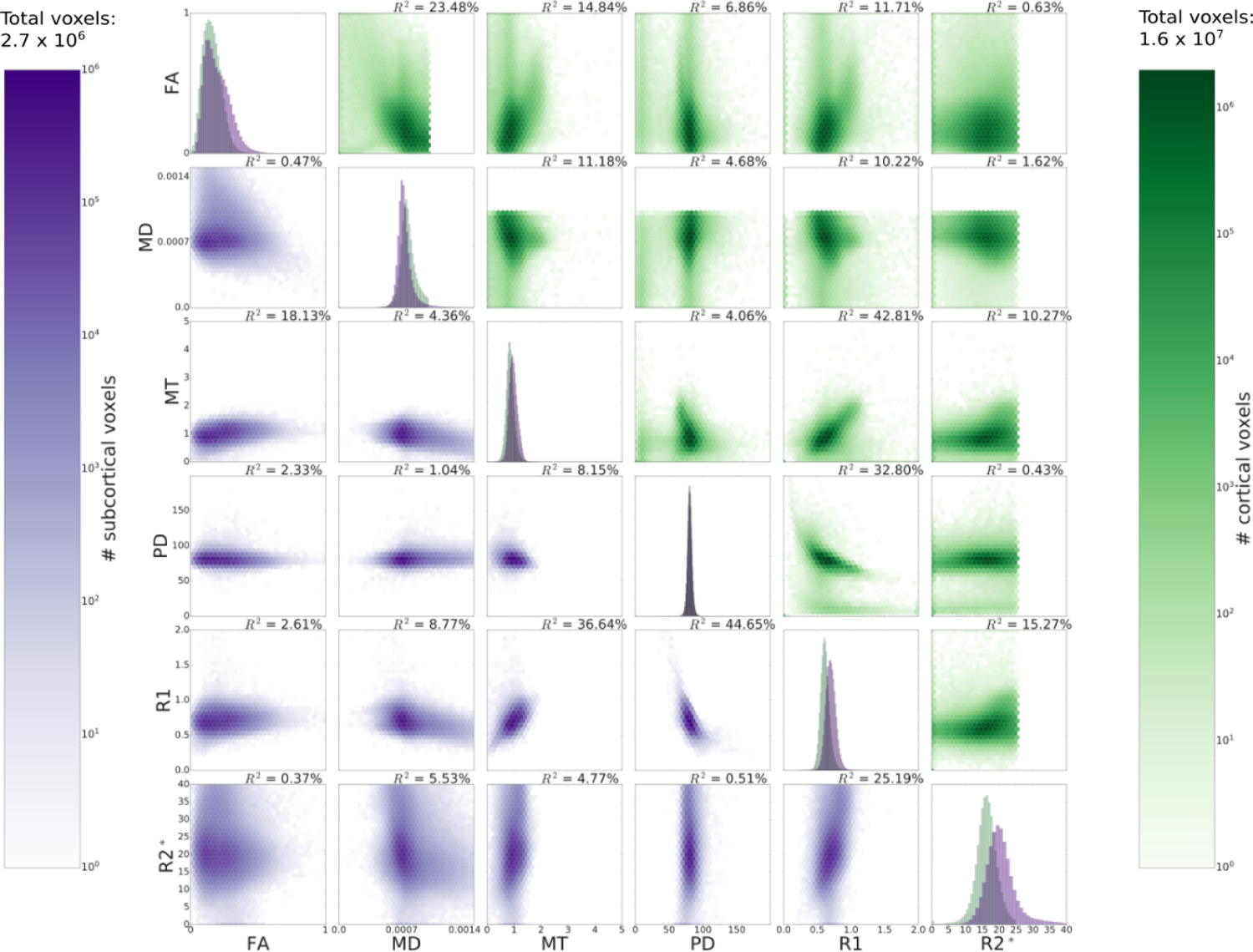
Pairwise correlations in cortical GM (above the diagonal) and subcortical GM (below the diagonal) for all participants’ data. Data along the diagonal shows the histograms for each metric in cortical (green) and subcortical GM (purple). Metric thresholds were implemented for R2* and MD to reduce partial volume effects and artifacts, and are visible as sharp cutoffs in the figure (see Method). Pairwise density plots are depicted in log space to show the full range of data.

### 3.4 Multimodal data visualization through dimensionality reduction

The results of the PCA revealed various patterns of cortical gradients. These analyses were done on all participants, but were also stable when applied to individuals, as suggested by the low variance in correlations across participants (Supplementary Figure 3 & 4). All PCA components, when mapped back onto the cortex, provided informative gradients, often denoting primary sensory and motor areas (Figure 4). Interestingly, the components that explained the most variance in the metrics seemed to diverge the least from the gradients of the metrics themselves. For example, PC1, which explains the most variance (44.0%), had a high weighting on R1 and metrics with strong correlations with R1. PC1’s cortical gradient showed high correspondence with primary sensory and motor regions, similar to the R1 cortical map (Figure 1). Similarly, PC2 resembled an inverse mapping of R2* and FA gradients, while PC3 resembled PD (see Supplementary Figure 2). The most novel gradient was in the component that explained the least amount of variance in contrasts (PC6, 3.1%) (Figure 4). This gradient least resembled individual quantitative metrics and showed a structural pattern having a degree of resemblance with the Default Mode Network, with high values in the posterior cingulate cortex, medial PFC, and posterior parietal cortex [Huntenburg et al., 2017; Raichle, 2015].

**Figure 4:**
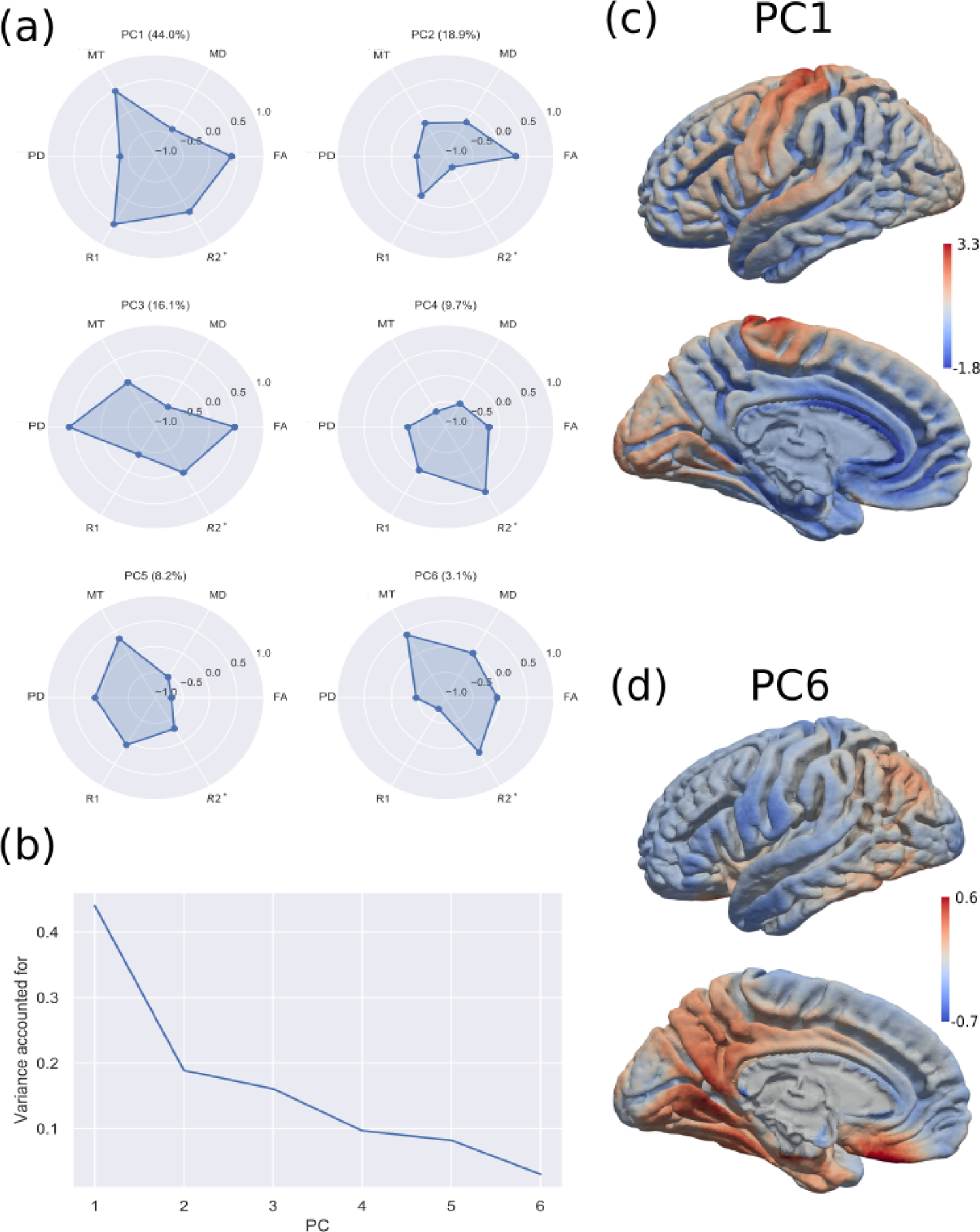
Overview of main results of the PCA. (a) radar plots showing the weighting of each component (ranging from −1 to 1) on each PC. In brackets is the percent variance accounted for by individual PCs. (b) A plot of variance accounted for in each PC. (c) PC1 mapped back onto the brain, showing primary sensory and motor regions. (d) PC6 mapped back onto the brain. Although this only accounts for 3.1% of the variance in the data, this component shows an interesting cortical gradient.

## Discussion

The goal of this study was to assess a set of quantitative metrics to provide a statistical description of their relationships. We used a set of previously established methods for extracting and co-registering the cortical ribbon [Bazin et al., 2014; Tardif et al., 2015] to identify core GM voxels that were less likely to suffer from partial volume effects. Bivariate correlations between metrics were performed for each tissue type, and our results showed that quantitative metrics have wide variations in their degree of correlation with each other, but are consistent across participants. Furthermore, we found that applying PCA resulted in the extraction of latent space components not obvious in individual unimodal metrics in the component accounting for the least variance.

Firstly, we observed the strongest overall pairwise correlation between R1 and MT, and the strength of this correlation was highly stable across tissue types (R^2^ = ∼36-48%). This may be due to the strong link between R1 and myelin [Möller et al., 2019; Trampel et al., 2019; Weiskopf et al., 2015] and MT as a measure of macromolecular tissue content [Filo et al., 2019; Henkelman et al., 2001; van der Weijden et al., 2021; Yarnykh et al., 2018]; both of which are expected to be highly related throughout the brain [Callaghan et al., 2015a; Khodanovich et al., 2017]. The normative R1-MT relationships described here could potentially be affected by pathological conditions that affect macromolecular proportions, such as Alzheimer’s tau pathologies and white matter hyperintensities, and baseline relationships would then be useful for research in these domains [van Es et al., 2006; Hentschel et al., 2004; Wardlaw et al., 2016].

Conversely, R2* had, on average, low correlations with other metrics (R^2^ average ∼= 7%), indicating little redundancy. Its strongest correlation was with R1 (GM R^2^ ∼= 21%, WM R^2^ ∼= 14%) which is consistent with previous research indicating a link between R2* and myeloarchitecture [Mangeat et al., 2015; Stüber et al., 2014], and an indirect link to myeloarchitecture through its relation with tissue iron concentration [Kirilina et al., 2020; Möller et al., 2019; Trampel et al., 2019; Weiskopf et al., 2015]. Interestingly, the R1-R2* correlation was weaker in WM where we might expect it to be highest based on their shared link with myeloarchitecture. One potential reason for this is that R2* is known to be affected by white matter fiber orientation relative to the main magnetic field (B0) [Kor et al., 2019; de Pasquale et al., 2013]. This orientation-dependent effect could lead to variation in R2* that does not arise from myeloarchitecture or iron and therefore decreases the strength of the correlation within the white matter. This effect is not expected to be significant within grey matter since there is no predominant net orientation of fibers at voxel resolutions. Overall, the differences in R1 and R2* correlations provide further support for the hypothesis that R1 is more strongly related to other aspects of microstructure, such as the cyto- and myeloarchitecture [Eickhoff et al., 2005] than R2*. Interestingly, the greater R1-R2* correlation in GM was predominantly driven by subcortical GM. Subcortical nuclei are relatively rich in iron [Drayer et al., 1986] and we collapsed across all identified subcortical GM structures (caudate, putamen, thalamus, and globus pallidus), which may lead to the greater variability seen in the subcortex than cortex for both metrics (as indicated by the spread in histograms along the diagonal of Figure 3). This greater variability may in turn drive the greater R1-R2* correlation by making it easier to identify a statistical correlation. This leads us to expect that changes in iron distribution throughout the brain and particularly in the basal ganglia, as is often seen in neurodegeneration and related pathologies [Carver, 2019; Daugherty and Raz, 2013; Khattar et al., 2021], may exhibit different R1-R2* correlations than what we have observed. Future work could seek to link the relationship between R1-R2* with other biomarkers of myelination and iron in patients diagnosed with pathologies related to iron accumulation for use as a non-invasive marker.

dMRI metrics (FA, MD) showed higher correlations with other metrics in GM than WM, as well as higher correlations in cortical GM than subcortical GM. This suggests that dMRI metrics may be dependent on tissue parameters that vary more independently in WM such as fiber orientation and directional uniformity, as would be expected from the diffusion tensor [Pierpaoli and Basser, 1996]. This then suggests that dMRI metrics in GM are more strongly driven by non-axonal microstructural differences than in white matter. For example, it is possible that differences in axonal radius and volume, as well as cell types and shapes could be a source of this change in dMRI correlations between the cortical and subcortical GM [Szafer et al., 1995; Szafer et al., 1995]. While FA differences/changes have previously been inferred to be due to differences in myelination, fiber orientation, and/or density [Beaulieu, 2014; Concha et al., 2010; Zatorre et al., 2012], we found that there were low correlations between FA and R1 (GM R^2^ ∼ 13%; WM R^2^ ∼ 8%; cortical R^2^ ∼ 12%; subcortical R^2^ ∼ 3%), which is arguably a more direct indicator of myelin [Trampel et al., 2019]. Past research has found that R1 and diffusion have differential variation throughout the lifespan, further supporting a difference in the information captured by these metrics, and suggesting that varying R1-dMRI correlations may be informative about health and ageing related issues [Yeatman et al., 2014]. The relatively low correlation in WM is also likely to be partially driven by the mismatch between the tensor-based simplification of WM architecture as FA, which best represents a single fibre population, and the reality that most regions include multiple fibre populations [Figley et al., 2022]. It may be necessary to combine more fibre-specific parameters from multishell DWI [Raffelt et al., 2017] with fibre-specific T1/R1 estimates from inversion-recovery DWI [De Santis et al., 2016; Leppert et al., 2021] to more accurately specify this relationship. Our findings are in line with decades of work characterizing the relationships between MR metrics in WM and support the idea that differences/changes in FA may not generally be attributable to a specific anatomical or physiological property such as myelin [Geeraert et al., 2019; Jones et al., 2013].

The correlation between R1 and PD had a stronger correlation in GM than WM, and this effect was most prominent in subcortical GM, similarly to the R1-R2* relationship. We do not have a clear interpretation for this R1-PD relationship. However, previous literature suggests that PD and T1/R1 are related through their dependence on cell densities - which affects free water content [Tardif et al., 2016]. Furthermore, the echo averaging prior to the map calculation introduces an R2* bias on the PD estimates (Weiskopf et al., 2013), which may be a partial driver of the stronger correlation of R1 and PD in subcortical GM.

We were also interested in exploring how dimensionality reduction could be used to determine whether combining metrics enhances details that are difficult or impossible to detect with single metrics [Geeraert et al., 2019]. Consistent with the pattern that we identified in the bivariate statistics, R1’s high covariance with multiple metrics appeared in the PCA as a cluster of metrics defining PC1 (including high loadings on all metrics except R2* and accounting for 44% of the variance). This component could be a potential candidate for a more specific index of cortical myelination [Lutti et al., 2014; Sereno et al., 2013]. PC1, along with most other PCs, resembled gradients found in individual metrics when mapped onto the cortex, such as that of primary sensory and motor cortices seen in R1. However, the last PC, which accounted for ∼3% of the variance in the data, provided a gradient containing some regions similar to those in the default-mode network (DMN) commonly identified in functional resting-state MRI (Raichle, 2015; Yeo et al., 2011). Previous research has shown links between the DMN and structural connectivity exhibited by diffusion tensor imaging and tractography [Horn et al., 2014], and between functional connectivity and cortical qT1 as a proxy for intracortical myelin [Huntenburg et al., 2017]. Following from this work, it would be interesting to explore if the gradients exhibited by the PCs are more directly related to functional connectivity and the DMN than any single modality. Whether or not our last PC corresponds to the DMN, we believe that it is interesting because it was the most dissimilar PC to individual metric gradients, suggesting that such multimodal tools can uncover subtle gradients that are overshadowed by other sources of variation in the data [Geeraert et al., 2019].

### Future directions

One major limitation of the present study is that it provides only a description of the relationship between qMRI metrics, but cannot comment on their fundamental relationship with physiological tissue parameters. Properly addressing this question will require more studies combining qMRI sequences with ground truth histological and biomarker data in the same brains/tissues [Weiskopf et al., 2015; Weiskopf et al., 2021], although even these approaches are limited due to postmortem factors that alter microstrucure. While our study used linear dimensionality reduction to allow for clear interpretation, it is likely that proper mapping from MRI to histological variables will require non-linear statistical methods as well. An approach similar to the recently proposed Morphometric Similarity Networks [Seidlitz et al., 2018] could be applied to quantitative multiparametric imaging data to better specify structural covariance [Mechelli, 2005; Zielinski et al., 2010] and eventually link it to the underlying physiological parameters. In addition, future work with ultra-high resolution MRI could be used to investigate multiparametric differences in contrasts across cortical layers [Eickhoff et al., 2005; Marques et al., 2017; Tardif et al., 2015; Waehnert et al., 2016].

The current study describes the bivariate relationships between qMRI metrics and an initial exploration of how the data is grouped in multidimensional space. We found that there was a wide range of covariance between metrics that differed across tissue type, but that these relationships were stable across individuals. Our findings are a step towards using multimodal metric combinations to better identify tissue characteristics and develop a more direct link between non-invasive MRI and physiology.

## Supplementary Figures

**Supplementary Figure 1:**
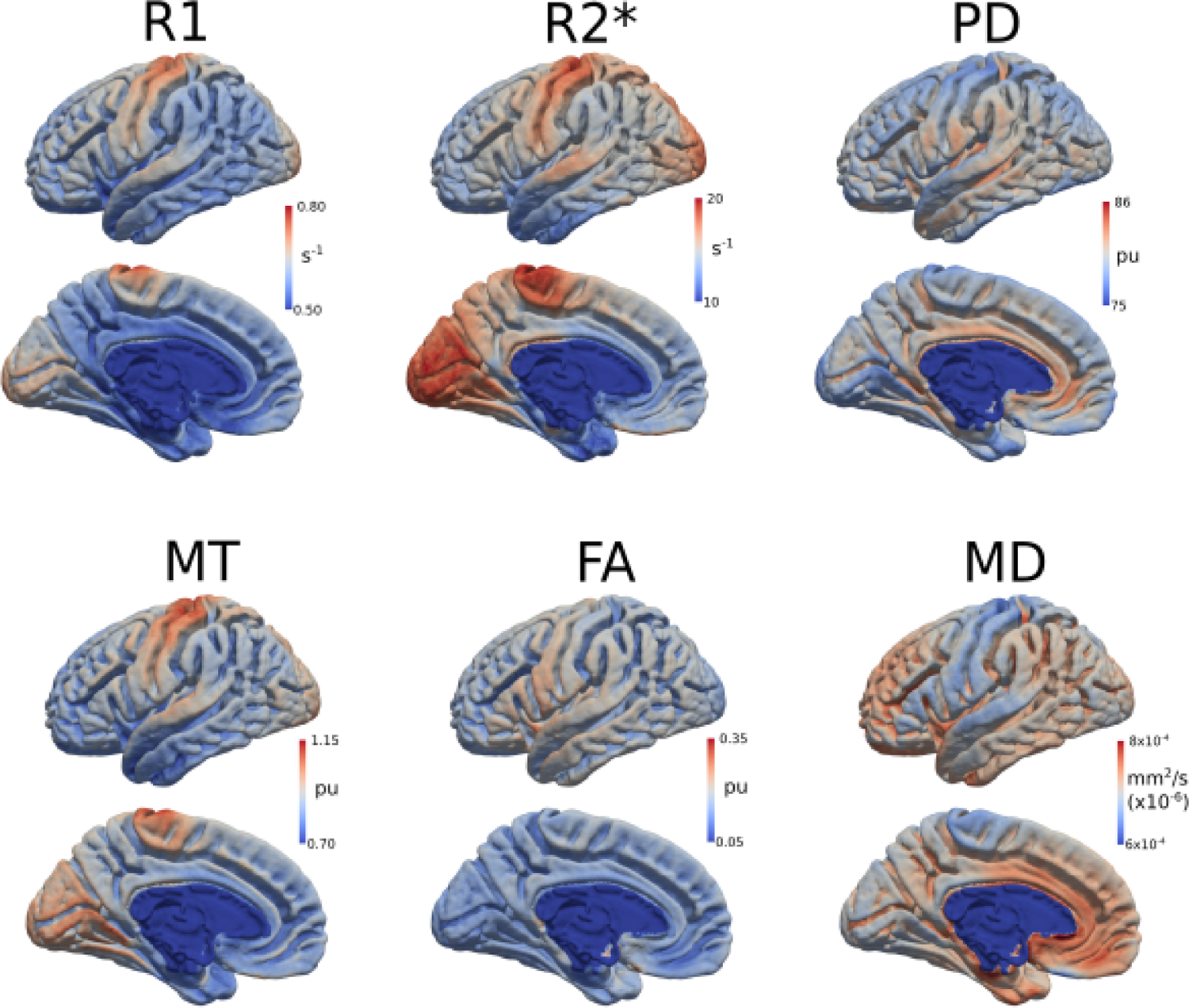
Surface plots for the six metrics. Median values from the cortical ribbon across all participants are plotted on the cortical surface. The range of values and units are described as well, with pu standing for percentage units.

**Supplementary Figure 2:**
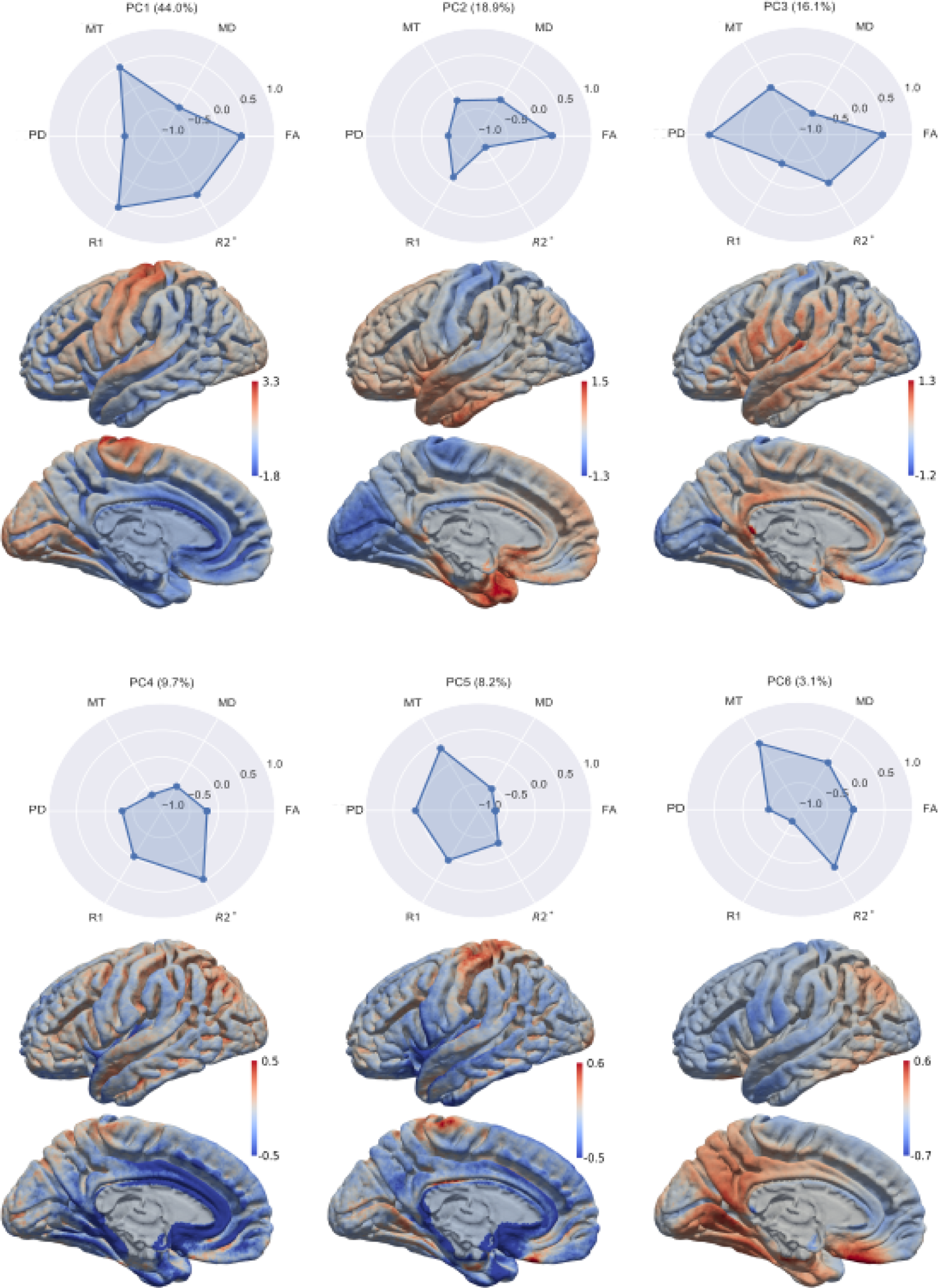
Surface plots and metric loadings for all PC’s. Median values across participants for each principal component was plotted on the cortical surface. The normalized loading of the metrics for each PC is also shown.

**Supplementary Figure 3:**
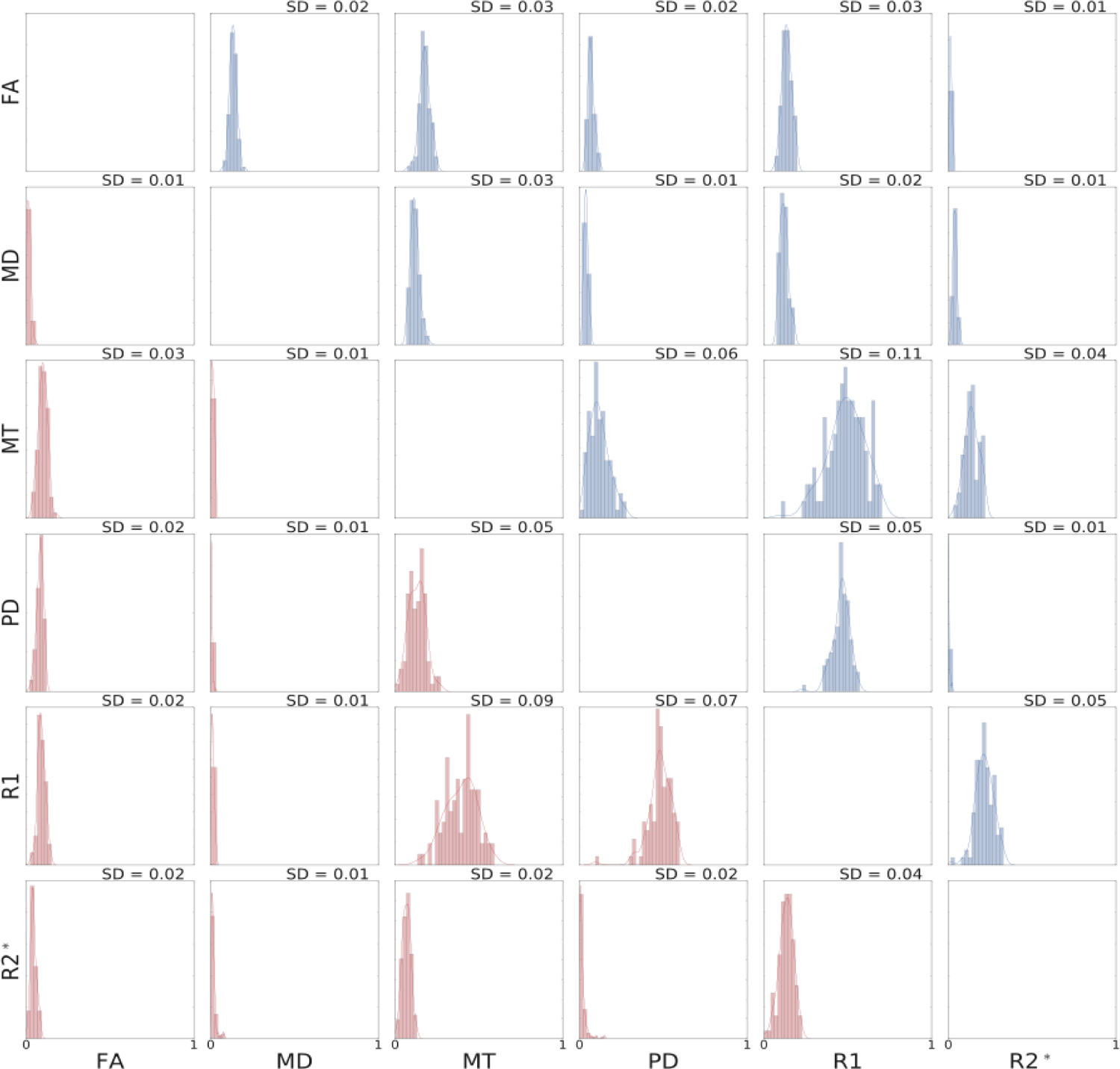
Distribution of R^2^ values across participants for all pairwise correlations (GM in blue, WM in red). This shows that the relationships between metrics were highly consistent across individuals.

**Supplementary Figure 4:**
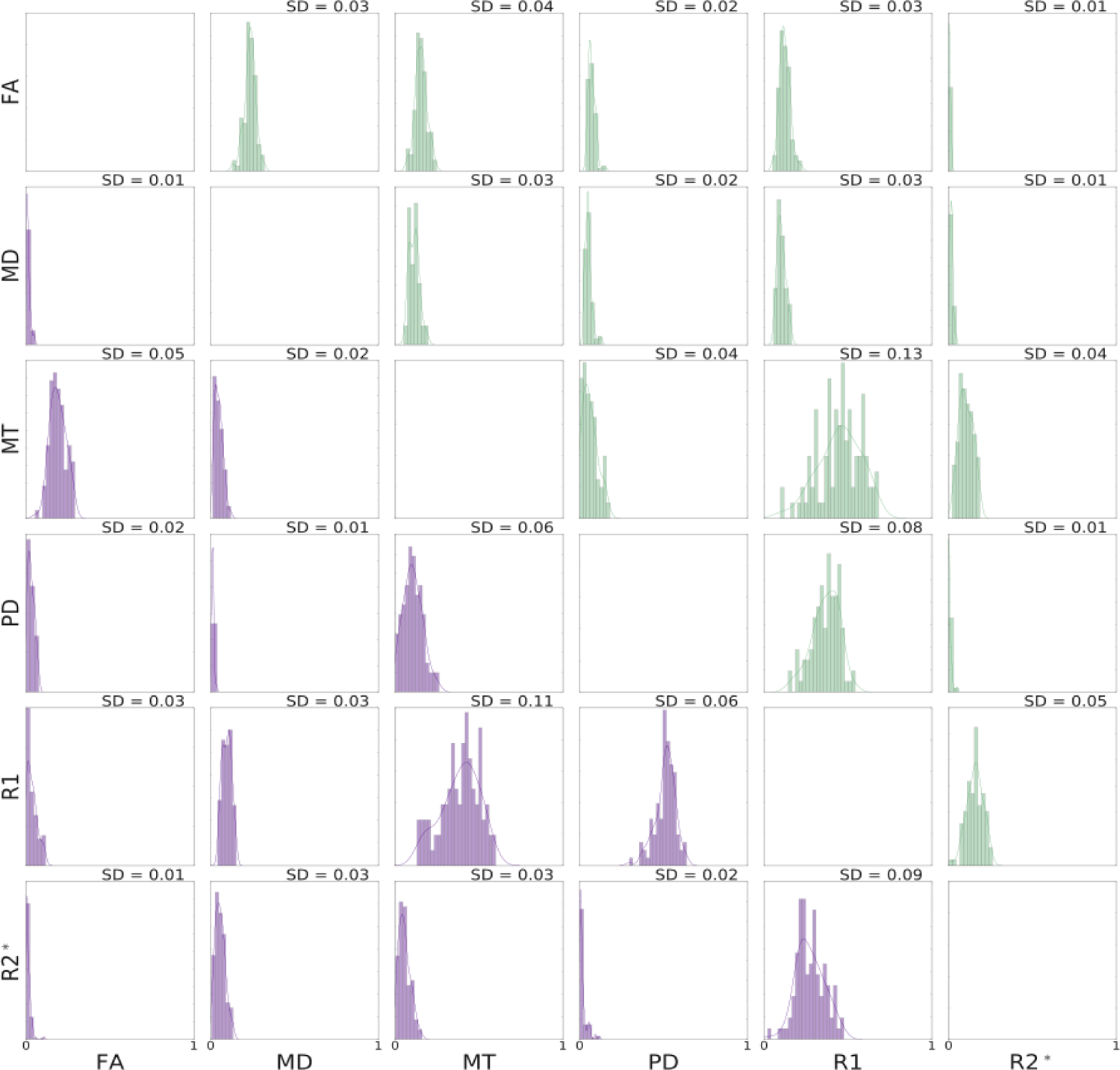
Distribution of R^2^ values across participants for all pairwise correlations (cortical GM in green, subcortical GM in indigo). This shows that the relationships between metrics were highly consistent across individuals.

## Acknowledgements

CJS was supported by the Natural Sciences and Engineering Research Council (NSERC: RGPIN-2020-06812, DGECR-2020-00146) and the Heart and Stroke Foundation of Canada New Investigator Award and Catalyst from the Canadian Institutes of Health Research (HNC 170723).

PLB was supported by the NWO Vici grant (PI: Birte Forstmann).

CJG was supported by the Heart and Stroke Foundation of Canada New Investigator Award, Michal and Renata Hornstein Chair in Cardiovascular Imaging, and NSERC (DG:

RGPIN-2015-04665).

AA and AF received funding from the SPP2041 program “Computational Connectomics” of the German Research Foundation (DFG).

NW has received funding from the European Research Council under the European Union’s Seventh Framework Programme (FP7/2007-2013) / ERC grant agreement n° 616905; from the European Union’s Horizon 2020 research and innovation programme under the grant agreement No 681094; from the BMBF (01EW1711A & B) in the framework of ERA-NET NEURON.

## Data and code availability statement

All code and parameters are available for public access in our github repository https://github.com/neuralabc/paper_QuantitativeMetricComparisons and summary maps will be made available on neurovault. Data is not available.

## References

1. Andersson JLR, Sotiropoulos SN (2016): An integrated approach to correction for off-resonance effects and subject movement in diffusion MR imaging. NeuroImage 125:1063–1078.

2. Ashburner J, Friston KJ (2005): Unified segmentation. NeuroImage 26:839–851.

3. Avants BB, Tustison NJ, Song G, Cook PA, Klein A, Gee JC (2011): A reproducible evaluation of ANTs similarity metric performance in brain image registration. NeuroImage 54:2033–2044.

4. Bazin P-L, Weiss M, Dinse J, Schäfer A, Trampel R, Turner R (2014): A computational framework for ultra-high resolution cortical segmentation at 7Tesla. NeuroImage 93:201–209.

5. Beaulieu C (2014): The Biological Basis of Diffusion Anisotropy. In:. Diffusion MRI. Elsevier. pp 155–183. https://linkinghub.elsevier.com/retrieve/pii/B9780123964601000081.

6. Bogovic JA, Prince JL, Bazin P-L (2013): A multiple object geometric deformable model for image segmentation. Comput Vis Image Underst 117:145–157.

7. Borsboom D, Mellenbergh GJ, van Heerden J (2003): The theoretical status of latent variables. Psychol Rev 110:203–219.

8. Caan MWA, Bazin P, Marques JP, Hollander G, Dumoulin SO, Zwaag W (2019): MP2RAGEME: T_1_, T_2_ *, and QSM mapping in one sequence at 7 tesla. Hum Brain Mapp 40:1786–1798.

9. Callaghan MF, Helms G, Lutti A, Mohammadi S, Weiskopf N (2015a): A general linear relaxometry model of R _1_ using imaging data: General Linear Relaxometry Model of R1. Magn Reson Med 73:1309–1314.

10. Callaghan MF, Josephs O, Herbst M, Zaitsev M, Todd N, Weiskopf N (2015b): An evaluation of prospective motion correction (PMC) for high resolution quantitative MRI. Front Neurosci 9. http://www.frontiersin.org/Brain_Imaging_Methods/10.3389/fnins.2015.00097/abstract.

11. Carver PL ed. (2019): 4. IRONING OUT THE BRAIN. In:. Essential Metals in Medicine: Therapeutic Use and Toxicity of Metal Ions in the Clinic. De Gruyter. pp 87–122. https://www.degruyter.com/document/doi/10.1515/9783110527872-004/html.

12. Concha L, Livy DJ, Beaulieu C, Wheatley BM, Gross DW (2010): In Vivo Diffusion Tensor Imaging and Histopathology of the Fimbria-Fornix in Temporal Lobe Epilepsy. J Neurosci 30:996–1002.

13. Dahnke R, Yotter RA, Gaser C (2013): Cortical thickness and central surface estimation. NeuroImage 65:336–348.

14. Daugherty A, Raz N (2013): Age-related differences in iron content of subcortical nuclei observed in vivo: A meta-analysis. NeuroImage 70:113–121.

15. De Santis S, Barazany D, Jones DK, Assaf Y (2016): Resolving relaxometry and diffusion properties within the same voxel in the presence of crossing fibres by combining inversion recovery and diffusion-weighted acquisitions: Resolving Myelin and Axonal Properties with IR-DTI. Magn Reson Med 75:372–380.

16. Draganski B, Ashburner J, Hutton C, Kherif F, Frackowiak RSJ, Helms G, Weiskopf N (2011): Regional specificity of MRI contrast parameter changes in normal ageing revealed by voxel-based quantification (VBQ). NeuroImage 55:1423–1434.

17. Drayer B, Burger P, Darwin R, Riederer S, Herfkens R, Johnson GA (1986): MRI of brain iron. Am J Roentgenol 147:103–110.

18. Eickhoff S, Walters NB, Schleicher A, Kril J, Egan GF, Zilles K, Watson JDG, Amunts K (2005): High-resolution MRI reflects myeloarchitecture and cytoarchitecture of human cerebral cortex. Hum Brain Mapp 24:206–215.

19. van Es ACGM, van der Flier WM, Admiraal-Behloul F, Olofsen H, Bollen ELEM, Middelkoop HAM, Weverling-Rijnsburger AWE, Westendorp RGJ, van Buchem MA (2006): Magnetization transfer imaging of gray and white matter in mild cognitive impairment and Alzheimer’s disease. Neurobiol Aging 27:1757–1762.

20. European Society of Radiology (ESR) (2015): Magnetic Resonance Fingerprinting - a promising new approach to obtain standardized imaging biomarkers from MRI. Insights Imaging 6:163–165.

21. Figley CR, Uddin MN, Wong K, Kornelsen J, Puig J, Figley TD (2022): Potential Pitfalls of Using Fractional Anisotropy, Axial Diffusivity, and Radial Diffusivity as Biomarkers of Cerebral White Matter Microstructure. Front Neurosci 15:799576.

22. Filo S, Shtangel O, Salamon N, Kol A, Weisinger B, Shifman S, Mezer AA (2019): Disentangling molecular alterations from water-content changes in the aging human brain using quantitative MRI. Nat Commun 10:3403.

23. Fukunaga M, Li T-Q, van Gelderen P, de Zwart JA, Shmueli K, Yao B, Lee J, Maric D, Aronova MA, Zhang G, Leapman RD, Schenck JF, Merkle H, Duyn JH (2010): Layer-specific variation of iron content in cerebral cortex as a source of MRI contrast. Proc Natl Acad Sci 107:3834–3839.

24. Geeraert BL, Lebel RM, Lebel C (2019): A multiparametric analysis of white matter maturation during late childhood and adolescence. Hum Brain Mapp 40:4345–4356.

25. Guilfoyle DN, Dyakin VV, O’Shea J, Pell GS, Helpern JA (2003): Quantitative measurements of proton spin-lattice (T1) and spin-spin (T2) relaxation times in the mouse brain at 7.0 T. Magn Reson Med 49:576–580.

26. Han X, Pham DL, Tosun D, Rettmann ME, Xu C, Prince JL (2004): CRUISE: Cortical reconstruction using implicit surface evolution. NeuroImage 23:997–1012.

27. Helms G, Dathe H, Kallenberg K, Dechent P (2008): High-resolution maps of magnetization transfer with inherent correction for RF inhomogeneity and *T* _1_ relaxation obtained from 3D FLASH MRI: Saturation and Relaxation in MT FLASH. Magn Reson Med 60:1396–1407.

28. Henkelman RM, Stanisz GJ, Graham SJ (2001): Magnetization transfer in MRI: a review. NMR Biomed 14:57–64.

29. Hentschel F, Kreis M, Damian M, Krumm B (2004): Leistet die Magnetization-Transfer-Ratio (MTR) einen Beitrag zur Diagnostik und Differenzialdiagnostik der Demenzen? RöFo - Fortschritte Auf Dem Geb Röntgenstrahlen Bildgeb Verfahr 176:1743–1749.

30. Horn A, Ostwald D, Reisert M, Blankenburg F (2014): The structural–functional connectome and the default mode network of the human brain. NeuroImage 102:142–151.

31. Huntenburg JM, Bazin P-L, Goulas A, Tardif CL, Villringer A, Margulies DS (2017): A Systematic Relationship Between Functional Connectivity and Intracortical Myelin in the Human Cerebral Cortex. Cereb Cortex 27:981–997.

32. Huntenburg JM, Steele CJ, Bazin P-L (2018): Nighres: processing tools for high-resolution neuroimaging. GigaScience 7. https://academic.oup.com/gigascience/article/doi/10.1093/gigascience/giy082/5049008.

33. Jenkinson M, Beckmann CF, Behrens TEJ, Woolrich MW, Smith SM (2012): FSL. NeuroImage 62:782–790.

34. Jones DK, Knösche TR, Turner R (2013): White matter integrity, fiber count, and other fallacies: The do’s and don’ts of diffusion MRI. NeuroImage 73:239–254.

35. Khattar N, Triebswetter C, Kiely M, Ferrucci L, Resnick SM, Spencer RG, Bouhrara M (2021): Investigation of the association between cerebral iron content and myelin content in normative aging using quantitative magnetic resonance neuroimaging. NeuroImage 239:118267.

36. Khodanovich MY, Sorokina IV, Glazacheva VY, Akulov AE, Nemirovich-Danchenko NM, Romashchenko AV, Tolstikova TG, Mustafina LR, Yarnykh VL (2017): Histological validation of fast macromolecular proton fraction mapping as a quantitative myelin imaging method in the cuprizone demyelination model. Sci Rep 7:46686.

37. Kirilina E, Helbling S, Morawski M, Pine K, Reimann K, Jankuhn S, Dinse J, Deistung A, Reichenbach JR, Trampel R, Geyer S, Müller L, Jakubowski N, Arendt T, Bazin P-L, Weiskopf N (2020): Superficial white matter imaging: Contrast mechanisms and whole-brain in vivo mapping. Sci Adv 6:eaaz9281.

38. Kor D, Birkl C, Ropele S, Doucette J, Xu T, Wiggermann V, Hernández-Torres E, Hametner S, Rauscher A (2019): The role of iron and myelin in orientation dependent R _2_ * of white matter. NMR Biomed:e4092.

39. Leppert IR, Andrews DA, Campbell JSW, Park DJ, Pike GB, Polimeni JR, Tardif CL (2021): Efficient whole-brain tract-specific T _1_ mapping at 3T with slice-shuffled inversion-recovery diffusion-weighted imaging. Magn Reson Med 86:738–753.

40. Leutritz T, Seif M, Helms G, Samson RS, Curt A, Freund P, Weiskopf N (2020): Multiparameter mapping of relaxation (R1, R2 *), proton density and magnetization transfer saturation at 3 T: A multicenter dual-vendor reproducibility and repeatability study. Hum Brain Mapp 41:4232–4247.

41. Lüsebrink F, Wollrab A, Speck O (2013): Cortical thickness determination of the human brain using high resolution 3T and 7T MRI data. NeuroImage 70:122–131.

42. Lutti A, Dick F, Sereno MI, Weiskopf N (2014): Using high-resolution quantitative mapping of R1 as an index of cortical myelination. NeuroImage 93:176–188.

43. Lutti A, Stadler J, Josephs O, Windischberger C, Speck O, Bernarding J, Hutton C, Weiskopf N (2012): Robust and Fast Whole Brain Mapping of the RF Transmit Field B1 at 7T. Ed. Wang Zhan. PLoS ONE 7:e32379.

44. Mancini M, Karakuzu A, Cohen-Adad J, Cercignani M, Nichols TE, Stikov N (2020): An interactive meta-analysis of MRI biomarkers of myelin. eLife 9:e61523.

45. Mangeat G, Govindarajan ST, Mainero C, Cohen-Adad J (2015): Multivariate combination of magnetization transfer, T 2 * and B0 orientation to study the myelo-architecture of the in vivo human cortex. NeuroImage 119:89–102.

46. Manning AP, Chang KL, MacKay AL, Michal CA (2017): The physical mechanism of “inhomogeneous” magnetization transfer MRI. J Magn Reson 274:125–136.

47. Margulies DS, Ghosh SS, Goulas A, Falkiewicz M, Huntenburg JM, Langs G, Bezgin G, Eickhoff SB, Castellanos FX, Petrides M, Jefferies E, Smallwood J (2016): Situating the default-mode network along a principal gradient of macroscale cortical organization. Proc Natl Acad Sci 113:12574–12579.

48. Marques JP, Khabipova D, Gruetter R (2017): Studying cyto and myeloarchitecture of the human cortex at ultra-high field with quantitative imaging: R1, R2* and magnetic susceptibility. NeuroImage 147:152–163.

49. Mechelli A (2005): Structural Covariance in the Human Cortex. J Neurosci 25:8303–8310.

50. Metere R, Kober T, Möller HE, Schäfer A (2017): Simultaneous Quantitative MRI Mapping of T1, T2* and Magnetic Susceptibility with Multi-Echo MP2RAGE. Ed. Vince Grolmusz. PLOS ONE 12:e0169265.

51. Möller HE, Bossoni L, Connor JR, Crichton RR, Does MD, Ward RJ, Zecca L, Zucca FA, Ronen I (2019): Iron, Myelin, and the Brain: Neuroimaging Meets Neurobiology. Trends Neurosci 42:384–401.

52. Paquola C, Vos De Wael R, Wagstyl K, Bethlehem RAI, Hong S-J, Seidlitz J, Bullmore ET, Evans AC, Misic B, Margulies DS, Smallwood J, Bernhardt BC (2019): Microstructural and functional gradients are increasingly dissociated in transmodal cortices. Ed. Henry Kennedy. PLOS Biol 17:e3000284.

53. de Pasquale F, Cherubini A, Péran P, Caltagirone C, Sabatini U (2013): Influence of white matter fiber orientation on R2* revealed by MRI segmentation. J Magn Reson Imaging 37:85–91.

54. Pedregosa F, Varoquaux G, Gramfort A, Michel V, Thirion B, Grisel O, Blondel M, Prettenhofer P, Weiss R, Dubourg V, Vanderplas J, Passos A, Cournapeau D Scikit-learn: Machine Learning in Python. Mach Learn PYTHON:6.

55. Pierpaoli C, Basser PJ (1996): Toward a quantitative assessment of diffusion anisotropy. Magn Reson Med 36:893–906.

56. Raffelt DA, Tournier J-D, Smith RE, Vaughan DN, Jackson G, Ridgway GR, Connelly A (2017): Investigating white matter fibre density and morphology using fixel-based analysis. NeuroImage 144:58–73.

57. Raichle ME (2015): The Brain’s Default Mode Network. Annu Rev Neurosci 38:433–447.

58. Roche A, Forbes F (2014): Partial Volume Estimation in Brain MRI Revisited:8.

59. Seidlitz J, Váša F, Shinn M, Romero-Garcia R, Whitaker KJ, Vértes PE, Wagstyl K, Kirkpatrick Reardon P, Clasen L, Liu S, Messinger A, Leopold DA, Fonagy P, Dolan RJ, Jones PB, Goodyer IM, Raznahan A, Bullmore ET (2018): Morphometric Similarity Networks Detect Microscale Cortical Organization and Predict Inter-Individual Cognitive Variation. Neuron 97:231–247.e7.

60. Sereno MI, Lutti A, Weiskopf N, Dick F (2013): Mapping the Human Cortical Surface by Combining Quantitative T1 with Retinotopy†. Cereb Cortex 23:2261–2268.

61. Shams Z, Norris DG, Marques JP (2019): A comparison of in vivo MRI based cortical myelin mapping using T1w/T2w and R1 mapping at 3T. Ed. Peter Lundberg. PLOS ONE 14:e0218089.

62. Stüber C, Morawski M, Schäfer A, Labadie C, Wähnert M, Leuze C, Streicher M, Barapatre N, Reimann K, Geyer S, Spemann D, Turner R (2014): Myelin and iron concentration in the human brain: A quantitative study of MRI contrast. NeuroImage 93:95–106.

63. Szafer A, Zhong J, Anderson AW, Gore JC (1995): Diffusion-weighted imaging in tissues: Theoretical models. NMR Biomed 8:289–296.

64. Tabelow K, Balteau E, Ashburner J, Callaghan MF, Draganski B, Helms G, Kherif F, Leutritz T, Lutti A, Phillips C, Reimer E, Ruthotto L, Seif M, Weiskopf N, Ziegler G, Mohammadi S (2019): hMRI – A toolbox for quantitative MRI in neuroscience and clinical research. NeuroImage 194:191–210.

65. Tardif CL, Gauthier CJ, Steele CJ, Bazin P-L, Schäfer A, Schaefer A, Turner R, Villringer A (2016): Advanced MRI techniques to improve our understanding of experience-induced neuroplasticity. NeuroImage 131:55–72.

66. Tardif CL, Schäfer A, Waehnert M, Dinse J, Turner R, Bazin P-L (2015): Multi-contrast multi-scale surface registration for improved alignment of cortical areas. NeuroImage 111:107–122.

67. Tardif CL, Steele CJ, Lampe L, Bazin P-L, Ragert P, Villringer A, Gauthier CJ (2017): Investigation of the confounding effects of vasculature and metabolism on computational anatomy studies. NeuroImage 149:233–243.

68. Trampel R, Bazin P-L, Pine K, Weiskopf N (2019): In-vivo magnetic resonance imaging (MRI) of laminae in the human cortex. NeuroImage 197:707–715.

69. Tustison NJ, Cook PA, Klein A, Song G, Das SR, Duda JT, Kandel BM, van Strien N, Stone JR, Gee JC, Avants BB (2014): Large-scale evaluation of ANTs and FreeSurfer cortical thickness measurements. NeuroImage 99:166–179.

70. Uddin MdN, Figley TD, Solar KG, Shatil AS, Figley CR (2019): Comparisons between multi-component myelin water fraction, T1w/T2w ratio, and diffusion tensor imaging measures in healthy human brain structures. Sci Rep 9:2500.

71. Waehnert MD, Dinse J, Schäfer A, Geyer S, Bazin P-L, Turner R, Tardif CL (2016): A subject-specific framework for in vivo myeloarchitectonic analysis using high resolution quantitative MRI. NeuroImage 125:94–107.

72. Wardlaw JM, Hernandez MCV, Munoz-Maniega S (2016): What are White Matter Hyperintensities Made of?: Relevance to Vascular Cognitive Impairment. J Am Heart Assoc 5. https://www.ahajournals.org/doi/10.1161/JAHA.115.002006.

73. van der Weijden CWJ, García DV, Borra RJH, Thurner P, Meilof JF, van Laar P-J, Dierckx RAJO, Gutmann IW, de Vries EFJ (2021): Myelin quantification with MRI: A systematic review of accuracy and reproducibility. NeuroImage 226:117561.

74. Weiskopf N, Edwards LJ, Helms G, Mohammadi S, Kirilina E (2021): Quantitative magnetic resonance imaging of brain anatomy and in vivo histology. Nat Rev Phys 3:570–588.

75. Weiskopf N, Mohammadi S, Lutti A, Callaghan MF (2015): Advances in MRI-based computational neuroanatomy: from morphometry to in-vivo histology. Curr Opin Neurol 28:313–322.

76. Weiskopf N, Suckling J, Williams G, Correia MM, Inkster B, Tait R, Ooi C, Bullmore ET, Lutti A (2013): Quantitative multi-parameter mapping of R1, PD*, MT, and R2* at 3T: a multi-center validation. Front Neurosci 7. http://journal.frontiersin.org/article/10.3389/fnins.2013.00095/abstract.

77. Yarnykh VL, Prihod’ko IY, Savelov AA, Korostyshevskaya AM (2018): Quantitative Assessment of Normal Fetal Brain Myelination Using Fast Macromolecular Proton Fraction Mapping. Am J Neuroradiol 39:1341–1348.

78. Yeatman JD, Wandell BA, Mezer AA (2014): Lifespan maturation and degeneration of human brain white matter. Nat Commun 5:4932.

79. Yeo BT, Krienen FM, Sepulcre J, Sabuncu MR, Lashkari D, Hollinshead M, Roffman JL, Smoller JW, Zöllei L, Polimeni JR, Fischl B, Liu H, Buckner RL (2011): The organization of the human cerebral cortex estimated by intrinsic functional connectivity. J Neurophysiol 106:1125–1165.

80. Zatorre RJ, Fields RD, Johansen-Berg H (2012): Plasticity in gray and white: neuroimaging changes in brain structure during learning. Nat Neurosci 15:528–536.

81. Zielinski BA, Gennatas ED, Zhou J, Seeley WW (2010): Network-level structural covariance in the developing brain. Proc Natl Acad Sci 107:18191–18196.

